# Temporal and spatial dynamics within the fungal microbiome of grape fermentation

**DOI:** 10.1101/2024.01.15.574562

**Authors:** Cristobal A. Onetto, Chris M. Ward, Steven Van Den Heuvel, Laura Hale, Kathleen Cuijvers, Anthony R. Borneman

## Abstract

Wine fermentation is a highly complex and competitive environment, imposing harsh selective pressures on fungal community ecology and diversity. The composition of fungal communities inhabiting the surface of grapes will directly impact fermentation progression, wine quality, and contribute to the distinctiveness between wines from different geographical regions. Despite this, the extent of microbial community diversity between geographies, termed ‘microbial terroir,’ remains highly debated. We amassed a large survey of grape spontaneous ferments over six years, encompassing 3105 fungal microbiomes across 14 geographically separated grape-growing regions, and nine grape cultivars. Investigation into the biodiversity of these ferments identified that few high abundance genera form the core of the initial grape microbiome. In line with previous studies, various consistent taxa were linked to specific geographical locations and grape varieties. However, these taxa accounted for a small portion of the overall diversity in the dataset. Through unsupervised clustering, we identified three distinct community types in the grape fungal microbiome, each exhibiting variations in the abundance of key genera. Analysing ferments across temporal and spatial scales revealed significant differences in species richness and compositional heterogeneity between wineries and grape growing regions. However, microbial communities were transient between years in the same winery, regularly transitioning between the three broad community types. We then investigated microbial community composition throughout the fermentative process and observed that initial microbial community composition is predictive of the diversity during the early stages of fermentation, with *Hanseniaspora uvarum* detected as the main non-*Saccharomyces* species within this large cohort of samples. Our results help to formulate a clear understanding of the spatial and temporal characteristics of the grape juice fungal microbiome and suggest that these communities are mainly defined by the grape niche and in a minor way shaped by local environmental conditions.

## Introduction

The surfaces of vines and grapes harbor a naturally occurring, intricate consortium of fungal and bacterial species, which exert a direct influence on vine growth and overall health [1-3]. These communities can vary significantly in their composition and may be affected by numerous factors, including grape variety, geographical location, climate and viticultural practices [4-6].

These microorganisms are transferred into the grape must during harvesting and processing, where they can play a pivotal role in shaping fermentation dynamics and contributing to the sensory attributes of the resulting wine. This influence is particularly significant during traditional uninoculated (“wild”) fermentation, where the absence of commercial yeast strains allows specific members of the microbial community to directly impact the fermentation process. Under this production framework, a complex microbial succession of yeasts and yeast-like fungi is generally observed as the grape must ferments. Initially, a diverse mixture of aerobic and apiculate yeasts, which normally reside on the surface of intact grape berries, dominate the fresh grape must. However, most of these species succumb to the decreasing oxygen levels and increasing ethanol concentrations, allowing mildly fermentative species to transiently increase in abundance [7-11]. Ultimately however, due to a higher fermentative ability, growth rate and tolerance to ethanol, most wild wine ferments converge to near monocultures of the major wine yeast, *Saccharomyces cerevisiae*. This is despite almost undetectable levels of this species being present on intact grape berries [12, 13].

While traditional microbiological techniques have provided important insights into the microbial succession that occurs in spontaneous ferments, both the breadth of ferments investigated and the depth at which individual species contributions could be resolved has been limited. Recent advances in culture-independent methods for species analysis, such as shotgun metagenomics and amplicon-based ITS phylotyping provides a high-throughput means to analyse large numbers of microbiological samples in great detail [14, 15].

Phylotyping techniques have been employed to investigate grape and fermentation microbiomes, aiming to uncover whether regional variations in wines can be partially attributed to the distinctiveness of microbial communities, supporting the notion of a “microbiome terroir”. While studies have revealed geographic patterns and considerable variation in microbial community composition among specific locations [4, 5, 16, 17], the extent to which these biogeographic patterns persist on a broader ecological scale remains uncertain. To address this, an extensive survey of grape juice samples was carried out to assess the microbial diversity inherent across a large cohort of uninoculated fermentations within Australia, that extended across nine grape varieties and 14 wine regions over the six successive vintages from 2016 to 2021. This comprehensive dataset enabled a thorough exploration of the core microbial communities in grape must and fermentation at a large ecological scale, thereby enhancing our understanding of the fungal species associated with grapes and the fermentation process.

## Materials and Methods

### Sample collection

A total of 36 wineries were sampled during six years across 14 different wine growing regions covering six states of Australia (Fig. 1a, Table S1). Wineries were supplied with pre-labelled, 50 mL falcon tubes. Approximately 40 mL of grape must was collected at four timepoints per fermentation tank, based upon refractometry levels (T1, freshly pressed grape juice; T2, after a 1 Bé drop in sugar; T3, 50% sugar remaining; T4, 10% sugar remaining) (Fig. 1b). Samples were stored frozen until collection and processing. All samples were obtained from strictly un-inoculated spontaneous ferments.

**Fig. 1:**
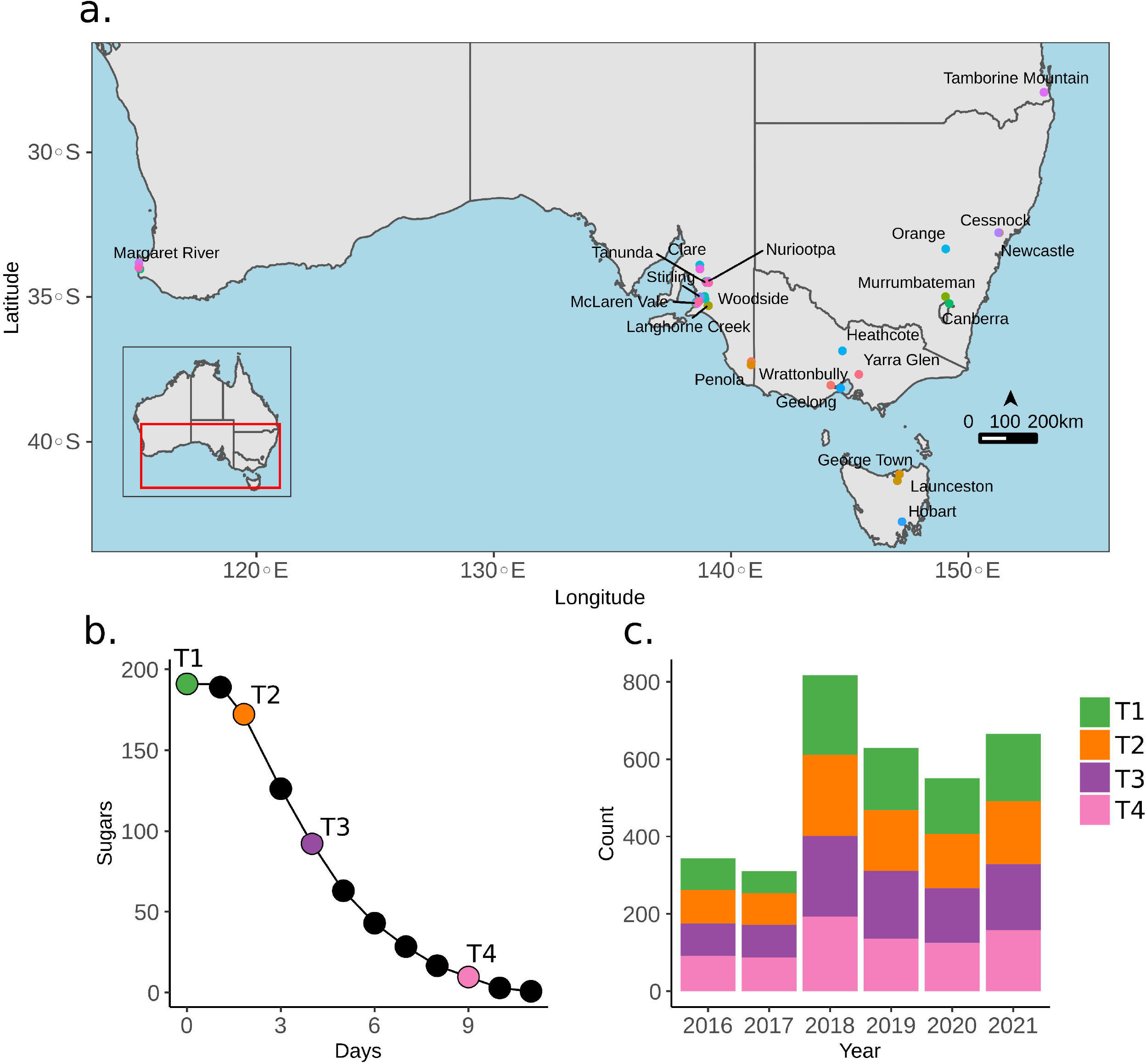
Sampling design of the study. **a** Geographical location of the Australian wineries sampled throughout a six-year period. Each point represents a winery and labels show the closest town/city. **b** Sampling point for each grape juice sample: T1= freshly pressed juice, T2= 1 Bé drop in sugar, T3= 50% sugar remaining; T4= 10% sugar remaining. **c** Number of samples collected each year for each timepoint.

### Sample processing and ITS sequencing

Samples were thawed, homogenized, and 1.9 mL of sample was centrifuged and washed twice with 1 mL of 1X PBS. DNA was extracted from pellets using the DNeasy PowerFood Microbial DNA Isolation Kit (Qiagen) following the manufacturer’s instructions. Bead-beating was carried out using a combination of 0.1 mm and 0.5 mm zirconia/silica beads (BioSpec Products) in a Precellys Evolution homogenizer (Bertin Instruments) at 8,000 RPM for 4 x 60 sec.

A two-step PCR was performed using 1 ng of DNA with primer sequences designed to amplify the fungal ITS region, while adding experiment-specific inline barcodes and appropriate adaptors for the Illumina sequencing platform as previously described by Sternes, Lee [14]. The first-round of amplification (20–30 cycles, 55°C annealing, 30-second extension, KAPA 2G Robust polymerase) targeting the ITS region was performed using primers BITS (ACCTGCGGARGGATCA) and B58S3 (GAGATCCRTTGYTRAAAGTT) [18]. 2 uL of first-round PCR product was used in the second amplification (15 cycles, 55°C annealing, 30-second extension, KAPA 2G Robust polymerase). Sequencing was performed using 2 x 300 bp paired-end chemistry in an Illumina instrument at the Ramaciotti Centre for Genomics (UNSW Sydney, Australia).

### Sequencing data processing

Raw sequencing reads were quality trimmed using Trimmomatic v0.38 [19], adaptor trimmed using Cutadapt v.1.16 [20] and merged using FLASH v2.2.00 [21]. Merged reads were hashed, de-replicated and counted as previously described by Sternes, Lee [14]. Operational taxonomic unit (OTU) clustering was performed using Swarm v2.2.2 [22]. Taxonomic classification was performed with the assign_taxonomy.py module of QIIME v1.9.1 [23] with a 98 % similarity cut off against the UNITE database (qiime_ver8_dynamic_10.05.2021). The taxonomic classification of all OTUs was also manually checked and curated against the BLAST database.

### Diversity analysis

All diversity analyses were performed in R v4.1.1 [24] and plots produced using a combination of ggplot2 [25] and the Ampvis2 package v2.8.3 [26]. Samples containing less than 2000 total read counts were removed from the dataset prior to analysis. Alpha (Shannon index) and beta diversity (Bray-Curtis dissimilarities) were calculated on rarefied OTU counts using Vegan v2.6 [27]. Differences of alpha diversity between categorical variables were investigated using Dunn’s multiple comparison test and Holm adjusted P-values. Investigation of differences in species diversity between categorical variables was tested using a permutational multivariate analysis of variance (MANOVA) [28] implemented in the adonis function with 999 permutations. Multivariate homogeneity of groups dispersions between categories was investigated using the betadisper function of Vegan and statistical differences between groups within categories were tested using a permutational ANOVA with 999 permutations. Pair-wise comparison between samples was performed with the permutest function implemented in Vegan. PERMANOVA is known to behave unreliably in unbalanced designs with heterogenous dispersion [29]. To address this issue the F_2_ statistic was [30] also used to test categories that showed heterogeneity in group dispersion (https://github.com/AWRI/permanovaModF). Investigation of specific taxonomic differences between categories was tested using the LDA effect size [31] on a normalised abundance table at genus level. Random forest supervised classification models [32] were used to evaluate the discriminative power of fungal with an LDA effect size >2. The strength of each predictor was evaluated using the mean decrease in Gini coefficient (MDG) as implemented in the R package Microeco [33], using 10000 trees per category with the randomForest package v.4.7 [34]. Community partitioning of T1 samples was performed using Dirichlet multinomial mixture (DMM) modelling [35] on a rarefied OTU abundance level matrix. The number of groups was obtained using the lowest Laplace estimation score. Microbiome branch structure was performed on the genus clustered relative abundance table using the PHATE algorithm v. 1.0.7 [36] with default parameters.

### Data availability

The sequencing data included in this study is available in NCBI under BioProject PRJNA1053885.

## Results

### The core fungal community of grapes is represented by few highly abundant species

To determine the core community of fungi associated with grapes and fermentation a large-scale survey of grape juice was conducted that covered 14 wine grape-growing regions of Australia (Fig. 1a). Samples were obtained from uninoculated grape juice samples across four timepoints during fermentation (Fig. 1b) from a total of 36 wineries throughout a period of six years (Fig. 1c), representing the largest microbial survey of grape juice to date. The fungal microbial community composition was characterized using amplicon sequencing targeting the ITS region. After quality filtering and OTU clustering, a total of 3105 samples were obtained containing 4309 OTUs.

T1 samples, which were taken as soon as fermentation tanks were filled with freshly crushed juice, largely reflected the composition of the fungal communities that inhabited the surface and flesh of grapes and were selected to estimate the grape-derived core fungal microbiome. While the T1 timepoint was shown to contain the highest diversity across the 772 samples, a small number of very abundant OTUs dominated the overall read abundance within most samples with the top 10 and 20 most abundant OTUs represented 70% (± 3.4%) and 80% (± 2.7%) of the total reads on average, respectively (Fig. 2a). OTUs that made up at least 80% of the reads in at least one sample were defined as abundant (Fig. 2b). To be considered a core OTU, we defined an abundance-occurrence threshold of 50% of the samples. While no single OTU was observed as abundant in all the samples investigated (Fig. 2b), two OTUs, one belonging to the genus *Aureobasidium* (observed in all samples, abundant in 659) and the other *Cladosporium* (observed in 770 samples, abundant in 424), passed the applied threshold for comprising core members of grape fungal microbiome (Fig. 2b). Species within the genus *Aureobasidium* are recognized for their ubiquitous presence in the phyllosphere of various plants, including grapevines [37] and these results confirm *Aureobasidium* as the predominant fungal genus across this large cohort of grape must samples (Fig. 2b and c). OTUs taxonomically assigned to *Cladosporium ramotenellum* (757 samples, abundant in 281), *Saccharomyces cerevisiae* (690 samples, abundant in 204) and *Hanseniaspora uvarum* (689 samples, abundant in 190) were also abundant across a large number of samples (Fig. 2b). When considering only the occurrence of individual OTUs, 54 OTUs were observed in at least 50% of the samples, 25 OTUs in 80% (Fig. 2c) and only a single OTU (genus *Aureobasidium*) in 100% of the samples analysed.

**Fig. 2:**
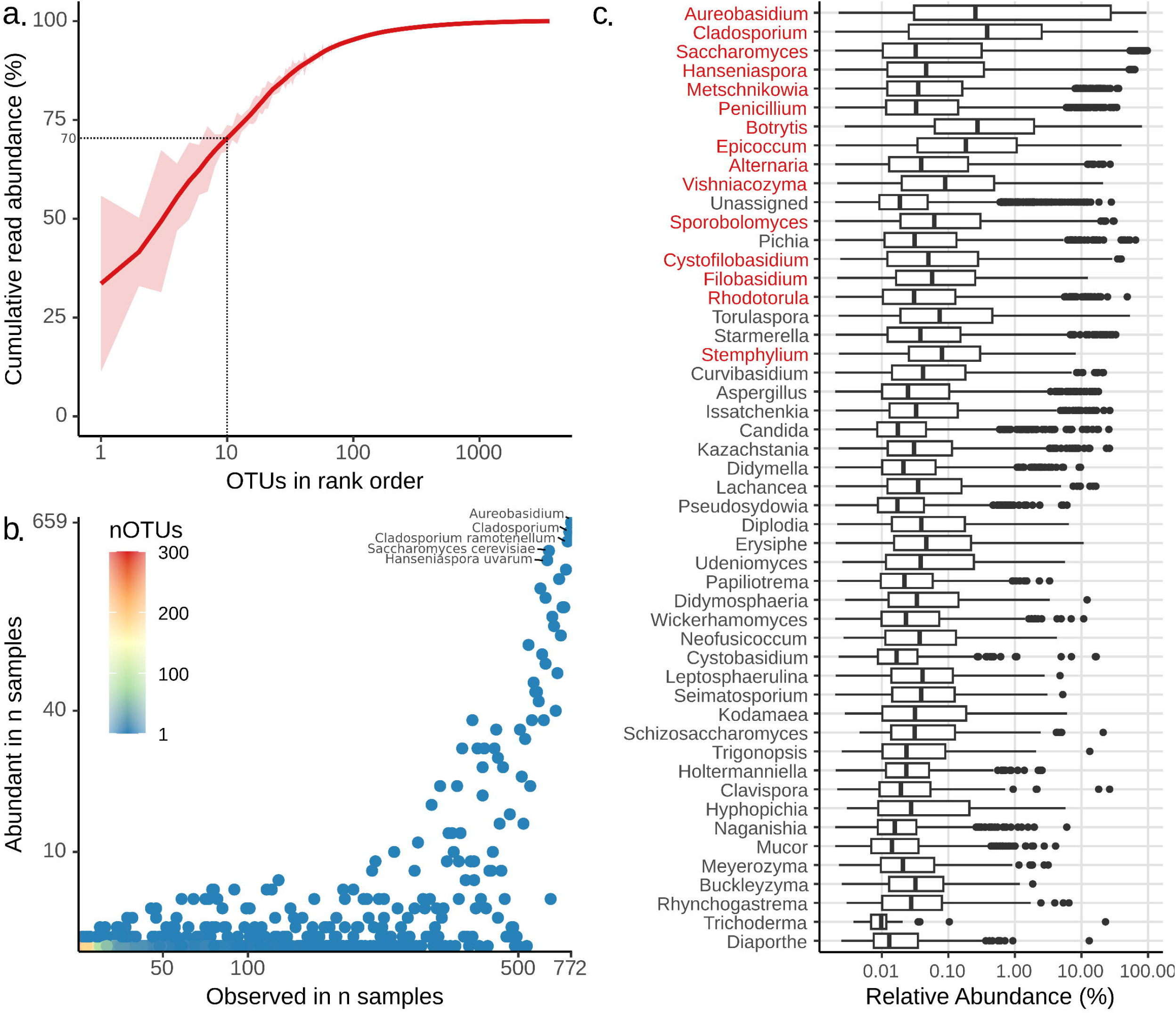
The core fungal community of grape juice. **a** Cumulative read abundance for 774 samples and 3594 OTUs. Red line represents the mean read abundance ± SD in red shading. OTUs were ranked in order of abundance for all samples. The 10 most abundant OTUs represent 70 % (± 3.4 %) of the total reads on average. **b** Frequency and observation as high abundant (OTUs making up the top 80% of the reads as in plot **a**) of OTUs across all samples. Points are coloured based on number of OTUs and Genus classification is shown for the top 5 OTUs observed as abundant. **c** Box plot showing the relative abundance of the top 50 genera sorted by mean read abundance. Genera in red represent taxa observed in at least 80% of all samples. Line inside the boxplot depicts the median and upper and lower ends of each box represent the 25th and 75th percentiles. Outer lines show the minimum and maximum value and dots represent potential outliers.

A global overview of all samples clustered by genus showed similar profiles to the OTU analysis and is consistent with previous studies [4, 6, 14, 38, 39]. The most abundant genera corresponded to *Aureobasidium, Cladosporium, Saccharomyces, Hanseniaspora* and *Metschnikowia* (Fig. 2c). As opposed to *Aureobasidium*, which comprised only four OTUs and with a single OTU accounting for most of the abundance (Fig. 2b), the genera *Cladosporium* (13 OTUs), *Saccharomyces* (67 OTUs), *Hanseniaspora* (46 OTUs) and *Metschnikowia* (53 OTUs) were comprised of high numbers of individual OTUs and suggests a higher species diversity within these genera.

### Temporal stability and regional variations in fungal communities

To investigate changes in fungal diversity over the course of the fermentation process, three sequential samples were collected after the T1 sampling point, as delineated by sugar consumption (Fig. 1b). The fermenting grape juice ecosystem exhibits a pronounced selective bias, favouring species with fermentative capabilities and increased tolerance to ethanol [40, 41]. Alpha and beta-diversity assessments across fermentation time points revealed clear reduction in both diversity and beta dispersion (Fig. 3a and Fig. S1a), accompanied with rapid shifts in microbial community composition (Fig. 3b) towards Saccharomycetes yeast species (Fig. S1b). In alignment with findings from many previous studies, 94% of T4 samples were predominantly dominated (> 50% relative abundance) by OTUs classified as *Saccharomyces cerevisiae*. Additionally, *Hanseniaspora uvarum, Schizozaccharomyces japonicus* and *Metschnikowia* were observed as the most abundant non-*Saccharomyces* at T4.

**Fig. 3:**
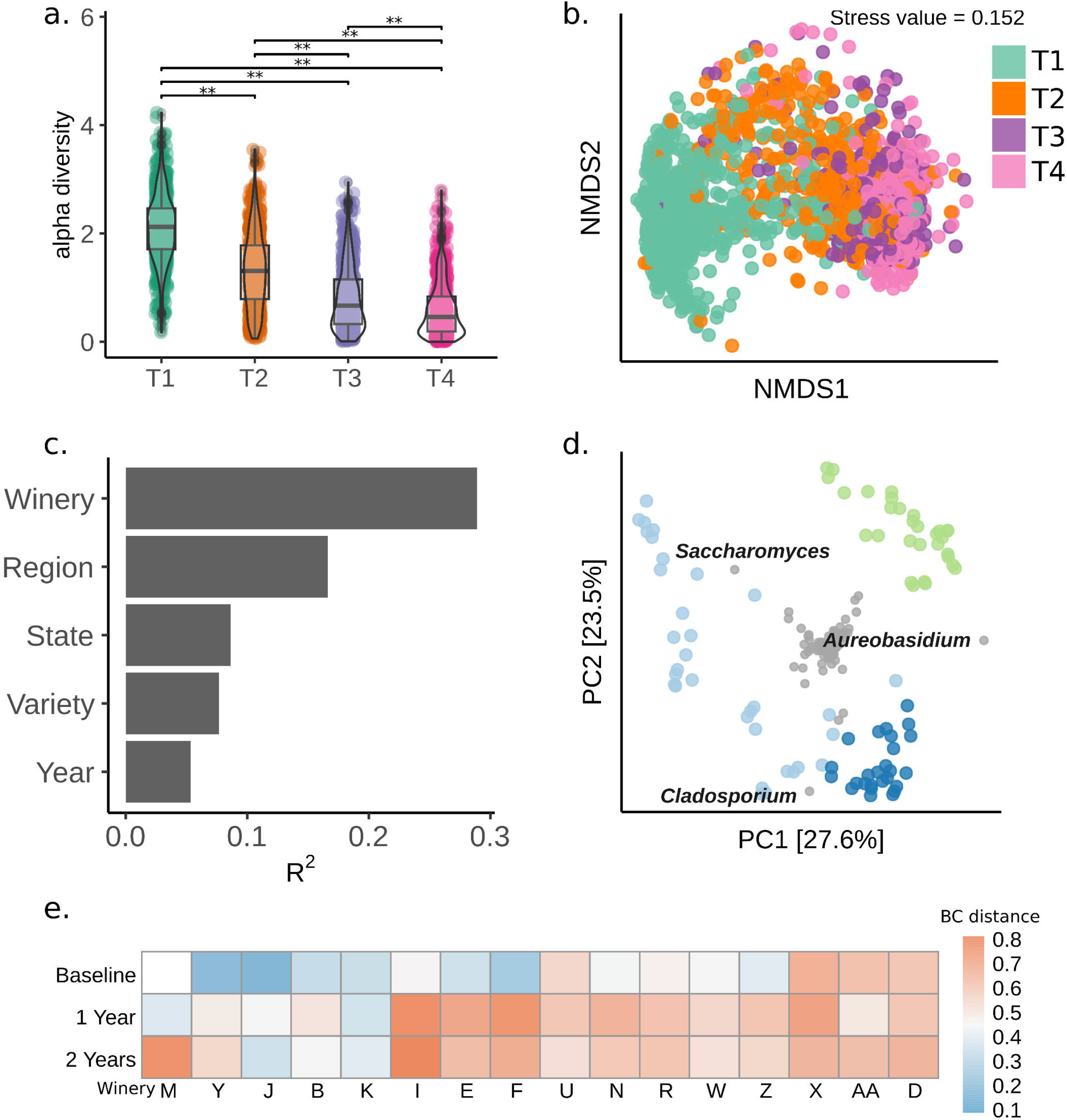
The fungal diversity of grape juice. **a** Violin-plot showing the alpha-diversity (Shannon index) of 3113 samples separated by sampling timepoint. Multiple-comparison test was performed between samples. **P⍰<⍰0.005 (Dunn’s, Holm-adjusted). **b** Non-metric multidimensional scaling (NMDS) of all samples based on Bray-Curtis (BC) dissimilarities. **c** Bar-plot showing the effect size (R^2^) of each metadata category calculated by PERMANOVA (Adonis). **d** Principal component analysis (PCA) of the Hellinger transformed OTU abundances of the three wineries displaying the highest F2 values after pairwise comparison (Table Sx). Individual OTUs are shown in grey with the three most extreme OTUs labelled to the genus level. **e** Temporal stability of the fungal communities. Baseline represents the average within BC distances for each winery in the year 2018. 1 and 2 years represent the verage BC distances between the baseline and 2019 and 2021, respectively.

Aside from *S. cerevisiae, H. uvarum* was observed as the most abundant yeast during the initiation of fermentation (1 Be drop), with 333 of 781 T2 samples containing greater than 10% relative abundance of H. uvarum. Additional genera of non-*Saccharomyces* species were also observed in high relative abundance (>10%) at the initiation of fermentation, albeit in a lower number of samples (*Aureobasidium*: 159, *Metschnikowia*: 77, *Cladosporium*: 50, *Pichia*: 29, *Issatchenkia*: 24, *Candida*: 18, *Starmerella*: 16, *Penicillium*: 15, *Torulaspora*: 14, *Schizosaccharomyces*: 7, *Botrytis*: 4 and *Lachancea*: 3 of 783 samples).

Significant differences in species richness (Dunn’s, Holm-adjusted p<0.05) and compositional heterogeneity (Permutest of beta dispersion p<0.05) were observed within T1 samples across various wineries and grape-growing regions (Fig. S2-S3, Tables S2-S5). Additionally, species richness and diversity exhibited substantial dispersion, indicating instability of the microbial communities (Fig. S2). To assess the temporal microbial stability, a comparative analysis of beta diversity was conducted across a panel of 16 wineries that were repeatedly sampled over three successive vintages (Fig. 3e). While a small number of wineries displayed a consistent microbial composition over time, the majority exhibited compositional alterations either after one or two years (Fig. 3e).

Previous studies have suggested that viticultural regions harbour regionally distinct microbial communities [4, 5, 16, 17]. To investigate this effect, a Permutational MANOVA of the Bray-Curtis dissimilarities between categories was utilized to assess fungal community structure between wineries, grape-growing regions, states, grape varieties, and years across the dataset (Fig. 3c). All tests were significant (Table S6), however a meaningful effect-size was only observed when comparing between wineries (Adonis R^2^ of wineries = 0.29) (Fig. 3c). To account for the observed heterogenous dispersion between categories, a modified F_2_ statistic was also calculated [30], producing similar results (Table S6). Principal Coordinates Analysis (PCoA) revealed no discernible clustering across the various categories examined (Fig. S4). Subsequent pairwise comparisons across individual wineries identified that the most pronounced dissimilarities were primarily associated with three specific wineries, namely K, E, and L (Table S7), situated within the Margaret River, Langhorne Creek, and Canberra District wine regions. Principal Component Analysis (PCA) analysis between these wineries indicated that they largely differ in the abundance of three OTUs corresponding to the genera *Aureobasidium, Cladosporium* and *Saccharomyces* (Fig. 3d).

To investigate significant associations between specific fungal genera and categories, a differential abundance testing method (LEfSE) combined with random forest models was implemented to evaluate the discriminative power of the differentially abundant genera. In agreement with previous studies [4, 5, 16, 17], significant associations (Log LDA score >2) were observed across all categories (Table S8 – S17). When focusing on broad geographical patterns, several genera were associated with specific states of Australia (Fig. 4a) with the largest MDG values observed for the genera *Cystofilobasidium* (SA), *Botrytis* (TAS) and *Sporobolomyces* (SA). Grape varieties also showed an influence on fungal microbiota (Fig. 4b) with *Dipodia* (Shiraz), *Seimatosporium* (Grenache) and *Cladosporium* (Chardonnay) amongst the top explaining taxa.

**Fig. 4:**
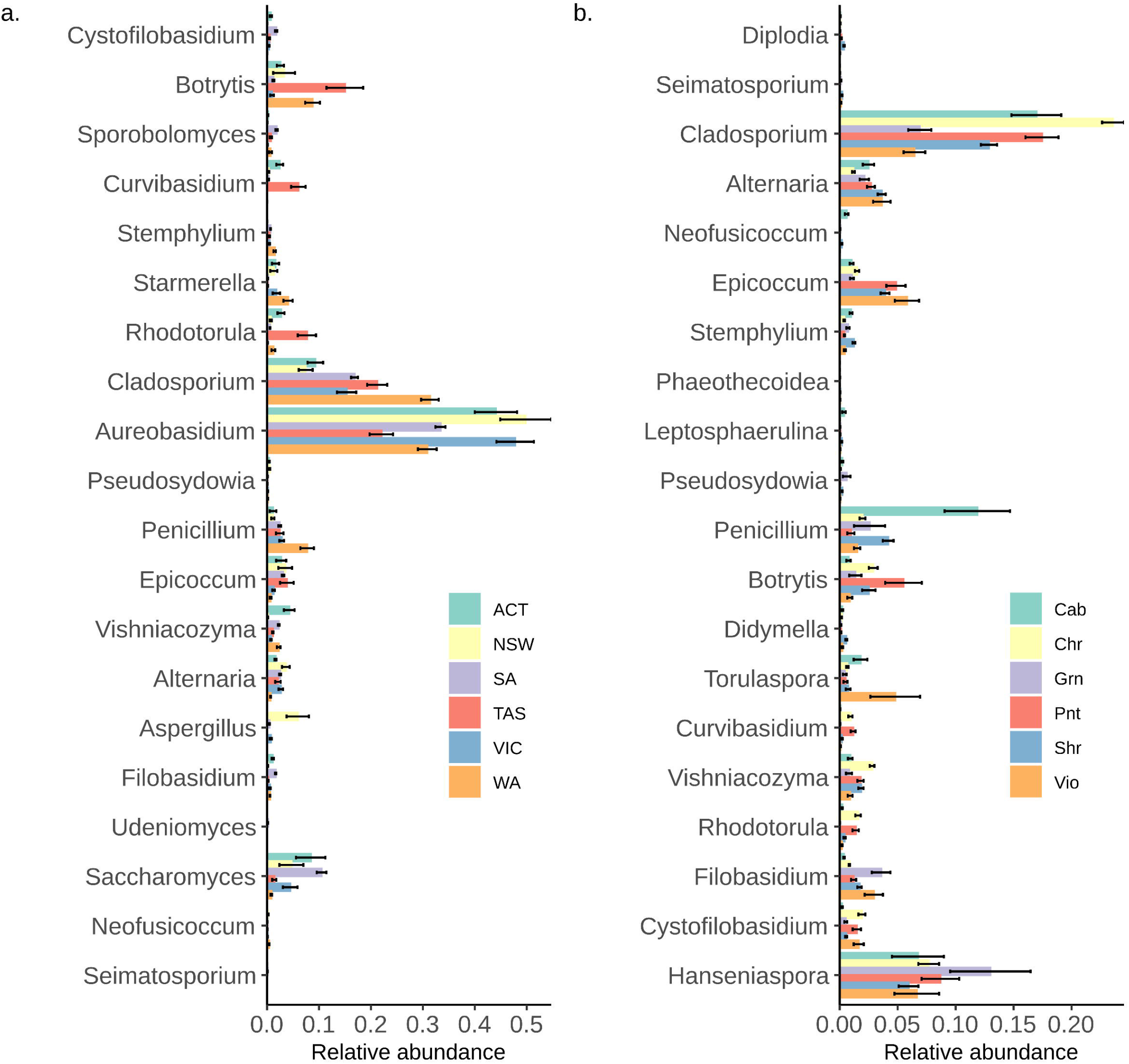
Taxa can distinguish between geographical regions and grape varieties. Relative abundance for the top 20 genera associated with **a** States of Australia and **b** grape varieties. Only taxa with a LDA effect size > 2 are shown. Taxa are sorted based on their Mean Decrease Gini of random forest models.

### Three community types partition the grape fungal microbiome

The large dataset provided the opportunity to investigate population stratification within the grape fungal microbiome and uncover microbial compositional patterns that would not be easily observed when focusing on individual taxa. To achieve this, a DMM-based approach [35], which has previously shown recurrent compositional patterns in multiple studies of the gut microbiome, was applied to this dataset [42].

Analysis of the 772 T1 samples indicated that the grape juice microbiomes could be partitioned into three distinct community types (defined as DMM1 - 3) (n DMM1 = 430, n DMM2 = 250 and n DMM3 = 92) that displayed clear ordination (Fig. 5a and b). Inspection of the 30 most abundant genera, in addition to the LefSE and random forest models, revealed clear differences between communities (Figure 5a, Table S18 – S19). DMM1 was defined by a high abundance of *Aureobasidium, Cladosporium, Metschnikowia* and several lower abundance genera (Fig. 5a). DMM2 displayed the highest abundance of *Aureobasidium* and the highest species richness (Fig. 5c), including the presence of several rare genera (Fig. 5a). DMM3 had the lowest species richness (Fig. 5c) and was mainly characterized by a high abundance of *Saccharomyces* and *Hanseniaspora* (Fig. 5a). Assignment of winery samples and wine growing regions to community types indicated that many were composed of mixed fungal community types (Fig. 5d).

**Fig. 5:**
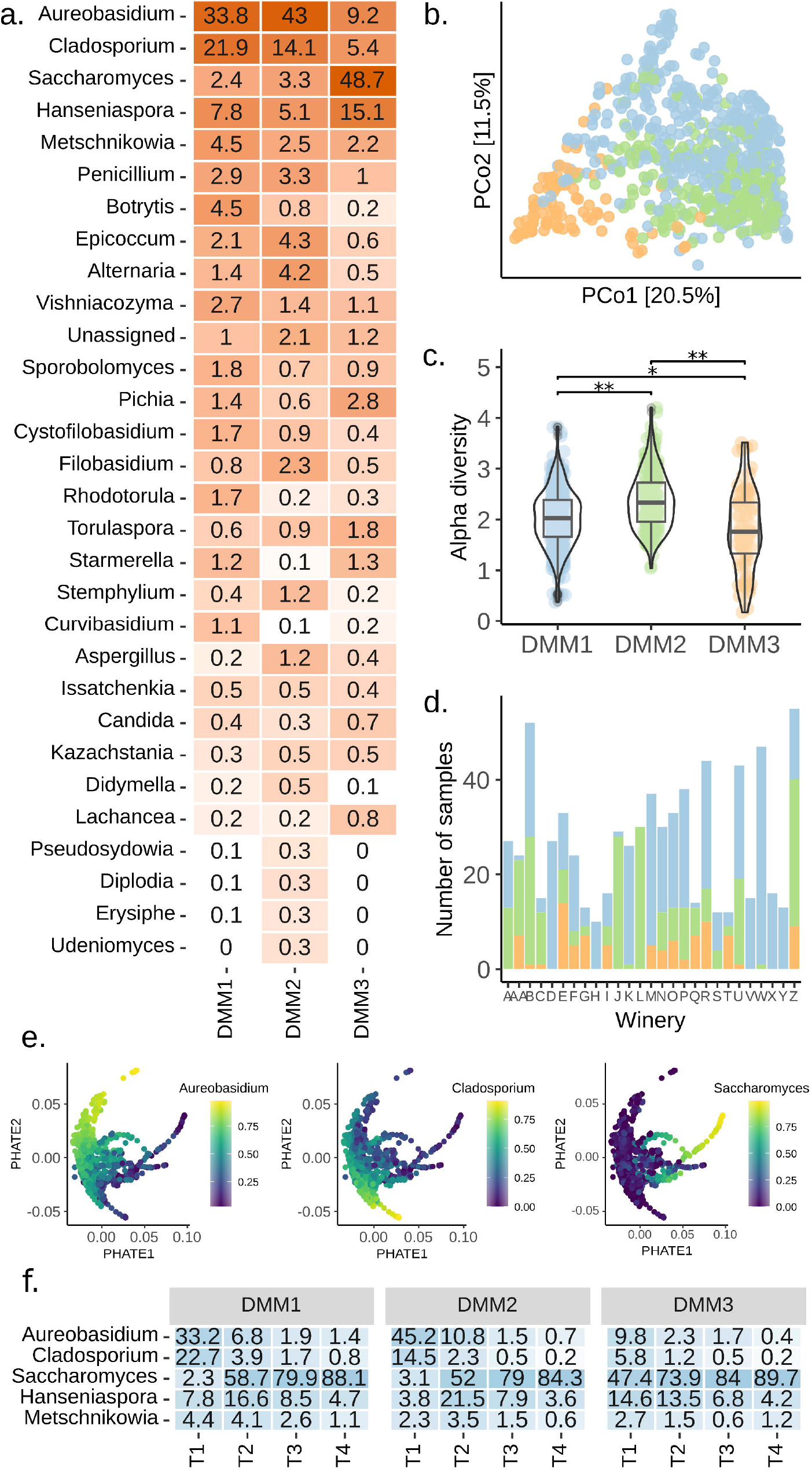
Dirichlet Multinomial Mixtures (DMM) partitioning of grape juice fungal communities. **a** Relative abundance for the top 30 Genera observed in each DMM partition. **b** Principal coordinate analysis (PCoA) of T1 samples based on Bray-Curtis (BC) dissimilarities. Samples are coloured by DMM as in panel (c) **c** Violin-plot showing the alpha-diversity (Shannon index) of samples belonging to each DMM partition. Multiple-comparison test was performed between samples. *P⍰<⍰0.05, **P⍰<⍰0.005 (Dunn’s, Holm-adjusted). **d** Total number of samples allocated to each DMM partition for each winery with more than 10 samples. Samples are coloured by DMM as in panel (c). **e** PHATE scatter plots where samples have been coloured based on the root squared transformed relative abundance of *Aureobasidium, Cladosporium and Saccharomyces*. **f** Relative abundance for the top 5 genera presented in panel (a) for timepoints T1 to T4 separated by DMM partition. Only DMM partition samples containing timepoints T2 to T4 were used to produce the plot.

It is possible that these three community types represent stages of ecological progression that are affected by external factors (e.g. grape maturity at sample time). To investigate this, the Potential of Heat-diffusion for Affinity-based Trajectory Embedding (PHATE) algorithm [36], a method developed to discover transitions in high dimensional data through the visualisation of branches and clusters was applied to the dataset. PHATE analysis showed three main branches (i.e. successions of microbiome arrangements bridging localized ecological conditions) that were associated with the abundance of the main genera discriminating between community types (Fig. 5e), suggesting transitions of microbial composition between these communities.

To evaluate the impact of initial community type on the microbial dynamics of fermentation, results from T2 through T4 were partitioned based on the community type assignment of their corresponding T1 sample. DMM1 and DMM2 retained an overall higher species richness at T2 (Fig. S1c) compared to DMM3. However, in all community types, fermentative species such as *Saccharomyces* and *Hanseniaspora* were already abundant at T2 (Fig. 5f). Non-fermentative taxa, namely *Aureobasidium* and *Cladosporium*, appeared to be more persistent in community types that initially contained a higher abundance of these genera in T1 (Fig. 5A). While this suggests that the initial community type directly influences the microbial dynamics of fermentation, it is evident that other external factors, such as grape juice composition and fermentation conditions, likely exert a substantial influence on microbial community dynamics.

## Discussion

The fungal communities of grapes have a direct effect on both the fermentation dynamics and the sensory attributes of spontaneously fermented wines [16, 43, 44]. To elucidate if the regional variation of wines could be partially explained by the distinctiveness of microbial communities, several studies have investigated the grape microbiota across different geographical scales using high-throughput amplicon sequencing, supporting the notion of a “microbiome terroir” [4, 5, 16, 17]. While it is clear from these studies that there can be extensive variation in microbial communities across geographical locations, they lack the sampling depth to obtain a global overview of the fungal communities of grape must. To address this, we carried out an extensive survey of 3105 grape juice samples across 14 grape-growing regions throughout a period of six years (Fig. 1), representing the most comprehensive microbial survey of grapes to date. Rather than concentrating on small scale regional or year-specific differences we analysed the dataset as a whole to elucidate general trends in community composition.

The core fungal community of grapes was surprisingly small, comprising only two OTUs and in addition, fewer than 20 OTUs accounted for the majority of the sequencing reads across the 772 T1 samples analysed. Relative to other open ecosystems, where the assembly of the communities is initiated by a multitude of species, grapes appear to display a very low number of core OTUs [45-49] and resemble the fungal microbiome of other cultivated fruit species [50]. Grapes therefore represent a highly selective ecosystem that differs significantly from other regions of the grapevine, such as the rhizosphere or bark, where a higher fungal taxonomic diversity have been observed [39, 51]. *Aureobasidium* appears to be particularly well adapted to the grape environment, with a single OTU dominating the abundance associated to this genus. While a single species could not be confidently assigned to this OTU, it is highly probable that it corresponds to *Aureobasidium pullulans*, given its more frequent isolation from grapes [52-55]. It is also interesting to note that community types featuring a higher abundance of this genus exhibit an overall greater species richness, suggesting that *Aureobasidium* species may not engage in broad antagonism but rather exhibit targeted antagonism against specific species, as reported in several studies [56, 57]. *Cladosporium*, known for causing post-harvest rot diseases in various fruits [58-60], is also part of the core microbiome of grapes, implying that its presence in grapes is not necessarily linked to disease development as previously suggested [39].

The fungal dynamics of fermentation progressed as expected [7, 8] with *S. cerevisiae* dominating by T2 in nearly all the samples analysed, yet many samples exhibited a prolonged period of high abundance of *Hanseniaspora uvarum*. Although prior research noted the prevalence of this species in early fermentation stages [61], these results highlight its role as the primary non-*Saccharomyces* yeast in grape juice fermentation. While other *Hanseniaspora* and non-*Saccharomyces* yeast were also detected at T1, their frequency was marginal by T2. From an adaptive viewpoint, *H. uvarum* seems less genetically equipped among Hanseniaspora species for fermentation due to a notable loss of genes related to cell-cycle and DNA repair [62]. Yet recent studies suggest that losing specific genes, including key cell-cycle regulators, might be an adaptive strategy, enabling a quicker cell cycle [63]. This discovery prompts the need for more research on *H. uvarum* to fully uncover its competitive advantage during early fermentation, especially compared to the various non-*Saccharomyces* yeast species found in grape must.

Wineries displayed distinct and temporally unstable fungal communities, and this is consistent with the observations reported in similar studies [4, 17, 39, 64, 65]. Despite this, the effect size of this associations was considerably lower than previously reported, likely due to differences in sampling size, geographic coverage, number of grape varieties and years of sampling. Large-scale geographic denominations exhibited significant associations with several fungal genera, however, the strongest variations between states and wineries were explained by species not considered to participate in grape fermentation and might reflect adaptations of the fungal microbiome to local environmental conditions, as previously suggested [4]. An example of this is the genus *Botrytis*, which was the second most important feature distinguishing the samples obtained from Tasmania, a geographical region known for having a colder climate and higher average rainfall, conditions that stimulate the growth of *Botrytis* species [66]. Several taxa distinguishing broader geographical regions coincide with microbiome reports from grape-growing areas in North America [4] and Europe [6]. Some of these genera include *Botrytis, Rhodotorula, Cladosporium, Aureobasidium*, and *Penicillium*. The congruence across studies provides support to the proposition by Bokulich [4] that biogeographical trends in different wine regions reflect responses to local conditions rather than patterns of microbial dispersion. Our results also suggests that grape variety plays a role in defining the abundance of select genera. Surprisingly, *Cladosporium* was observed as the top genus distinguishing the variety Chardonnay, which has been previously reported for grapes in North America [4], suggesting that either a genetic component or a standardised vineyard management approach associated with this variety drives the observed global pattern of microbial abundance. Variations in susceptibility to grapevine pathogens between varieties may also play a major role in explaining these differences. For example, *Diplodia* showed the highest MDG value for the variety Shiraz, which is known to be highly susceptible to the colonisation of the grapevine trunk pathogen *Diplodia serata* [67]. While our results support the proposition of a “microbial terroir”, the overlapping results with previous studies performed in different geographical regions suggest that global viticultural zones share a similar grape fungal microbiome that is mainly defined by the grape niche [1] and in a minor way shaped by local environmental conditions and grape variety.

Through an unsupervised partitioning of these large set of samples we were able to capture most of the microbial variability in three partitions that might reflect possible ecological states of the grape fungal microbiome. Rather than containing distinctive microbial communities, the data suggests that the disparities among geographical regions can be largely attributed to the prevalence of a few select genera, including *Aureobasidium, Cladosporium* and *Saccharomyces* and suggests that samples transition between three broad community types characterised by differences in diversity. While we do not know the factors driving the transitions between community types, previous reports have seen similar dynamics in the abundance of main genera (e.g. *Aureobasidium* and *Cladosporium*) throughout the developmental stages of grapes [39], suggesting that maturity of grapes at sampling time might be a driver between these transitions. These community types exhibited differences in several important genera known to impact wine composition (e.g., *Saccharomyces, Hanseniaspora, Metschnikowia, Pichia, Torulaspora, Starmerella* etc.) [68] and likely show functional heterogeneity with regards to fermentation. Indeed, more diverse communities retained a higher fungal diversity during the initial stages of fermentation, likely influencing fermentation performance and the chemical composition of wine. Therefore, future research should focus on understanding the environmental drivers behind shifts in community types, which could serve as a valuable tool to modulate grape health and fermentation.

## Conclusion

Taken together our results help to formulate a broader understanding of the community dynamics within grape fermentations across varietal, spatial and temporal scales utilizing the largest survey of grape ferment microbiomes to date. A holistic approach was taken with respect to the data analysis, investigating the role that geography, grape variety and year has on microbiome composition and diversity. When considering all scales, little structure was identified with wineries displaying highly unstable fungal communities that transitioned within three broad community types. In contrast to previous work suggesting that geography might play a large role in microbiome composition, it was observed that both regional and state level geographies showed a small effect size on the overall diversity and composition of ferments. In general, it was observed that initial community diversity was transient between fermentation timepoints while playing an important, but imperfect, role in predicting the diversity of the fungal microbiome throughout fermentation. This suggests that initial microbial community is not deterministic and highlights the role abiotic factors have in determining microbiome composition throughout the fermentation process.

## Acknowledgements

This work was supported by Australian grapegrowers and winemakers through their investment body Wine Australia, with matching funds from the Australian Government. The AWRI is a member of the Wine Innovation Cluster (WIC) in Adelaide.

## Data availability

The sequencing data included in this study is available in NCBI under BioProject PRJNA1053885.

## References

1. Morrison-Whittle P, Goddard MR. Quantifying the relative roles of selective and neutral processes in defining eukaryotic microbial communities. ISME J 2015;9:2003–2011. doi: 10.1038/ismej.2015.18.

2. Taylor MW, Tsai P, Anfang N, et al. Pyrosequencing reveals regional differences in fruit-associated fungal communities. Environ Microbiol 2014;16:2848–2858. doi: 10.1111/1462-2920.12456.

3. Müller DB, Vogel C, Bai Y, et al. The plant microbiota: systems-level insights and perspectives. Ann Rev Genet 2016;50:211–234. doi: 10.1146/annurev-genet-120215-034952.

4. Bokulich NA, Thorngate JH, Richardson PM, et al. Microbial biogeography of wine grapes is conditioned by cultivar, vintage, and climate. Proc Natl Acad Sci USA 2014;111:E139–E148. doi: doi:10.1073/pnas.1317377110.

5. Gayevskiy V, Goddard MR. Geographic delineations of yeast communities and populations associated with vines and wines in New Zealand. ISME J 2012;6:1281–1290. doi: 10.1038/ismej.2011.195.

6. Pinto C, Pinho D, Cardoso R, et al. Wine fermentation microbiome: a landscape from different Portuguese wine appellations. Front Microbiol 2015;6. doi: 10.3389/fmicb.2015.00905

7. Beltran G, Torija MJ, Novo M, et al. Analysis of yeast populations during alcoholic fermentation: A six year follow-up study. Syst Appl Microbiol 2002;25:287–293. doi: 10.1078/0723-2020-00097.

8. Combina M, Elía A, Mercado L, et al. Dynamics of indigenous yeast populations during spontaneous fermentation of wines from Mendoza, Argentina. Int J Food Microbiol 2005;99:237–243. doi: 10.1016/j.ijfoodmicro.2004.08.017.

9. Fleet GH. Growth of yeasts during wine fermentations. J Wine Res 1990;1:211–223. doi: 10.1080/09571269008717877.

10. Fleet GH. Wine yeasts for the future. FEMS Yeast Res 2008;8:979–995. doi: 10.1111/j.1567-1364.2008.00427.x.

11. Fleet GH, Lafon-Lafourcade S, Ribereau-Gayon P. Evolution of yeasts and lactic acid bacteria during fermentation and storage of bordeaux wines. Appl Env Microbiol 1984;48:1034–1038. doi: doi:10.1128/aem.48.5.1034-1038.1984.

12. Martini A, Ciani M, Scorzetti G. Direct enumeration and isolation of wine yeasts from grape surfaces. Am J Enol Viticult 1996;47:435–440. doi: 10.5344/ajev.1996.47.4.435.

13. Mortimer R, Polsinelli M. On the origins of wine yeast. Res Microbiol 1999;150:199–204. doi: 10.1016/S0923-2508(99)80036-9.

14. Sternes PR, Lee D, Kutyna DR, et al. A combined meta-barcoding and shotgun metagenomic analysis of spontaneous wine fermentation. GigaScience 2017;6:1–10. doi: 10.1093/gigascience/gix040.

15. Caporaso JG, Lauber CL, Walters WA, et al. Global patterns of 16S rRNA diversity at a depth of millions of sequences per sample. Proc Natl Acad Sci USA 2011;108:4516–4522. doi: 10.1073/pnas.1000080107.

16. Bokulich NA, Collins TS, Masarweh C, et al. Associations among wine grape microbiome, metabolome, and fermentation behavior suggest microbial contribution to regional wine characteristics. mBio 2016;7. doi:10.1128/mbio.00631-16.

17. Knight S, Klaere S, Fedrizzi B, et al. Regional microbial signatures positively correlate with differential wine phenotypes: evidence for a microbial aspect to terroir. Sci Rep 2015;5:14233. doi: 10.1038/srep14233.

18. Bokulich NA, Mills DA. Improved selection of internal transcribed spacer-specific primers enables quantitative, ultra-high-throughput profiling of fungal communities. Appl Environ Microbiol 2013;79:2519–2526. doi:10.1128/AEM.03870-12.

19. Bolger AM, Lohse M, Usadel B. Trimmomatic: a flexible trimmer for Illumina sequence data. Bioinformatics 2014:btu170.

20. Martin M. Cutadapt removes adapter sequences from high-throughput sequencing reads. EMBnet. Journal 2011;17:3. doi:10.14806/ej.17.1.200.

21. Magoč T, Salzberg SL. FLASH: Fast length adjustment of short reads to improve genome assemblies. Bioinformatics 2011. doi: 10.1093/bioinformatics/btr507.

22. Mahé F, Rognes T, Quince C, et al. Swarm: robust and fast clustering method for ampliconbased studies. PeerJ 2014;2:e593. doi: 10.7717/peerj.593.

23. Caporaso JG, Kuczynski J, Stombaugh J, et al. QIIME allows analysis of high-throughput community sequencing data. Nature methods 2010;7:335–336.

24. Team RC. R. A language and environment for statistical computing [Internet]. Vienna, Austria: R Foundation for Statistical Computing; 2013. Document freely available on the internet at: http://www.r-project.org2015.

25. Wickham H. ggplot2: elegant graphics for data analysis: Springer, 2016.

26. Skytte Andersen KS, Kirkegaard RH, Karst SM, et al. ampvis2: an R package to analyse and visualise 16S rRNA amplicon data. bioRxiv 2018:299537. doi: 10.1101/299537.

27. Dixon P. VEGAN, a package of R functions for community ecology. J Veg Sci 2003;14:927–930. doi: 10.1111/j.1654-1103.2003.tb02228.x.

28. Anderson MJ. A new method for non-parametric multivariate analysis of variance. Aust Ecol 2001;26:32–46. doi: 10.1111/j.1442-9993.2001.01070.pp.x.

29. Anderson MJ, Walsh DCI. PERMANOVA, ANOSIM, and the Mantel test in the face of heterogeneous dispersions: What null hypothesis are you testing? Ecol Monogr 2013;83:557–574. doi: 10.1890/12-2010.1.

30. Anderson MJ, Walsh DCI, Robert Clarke K, et al. Some solutions to the multivariate Behrens– Fisher problem for dissimilarity-based analyses. Aust NZ J Stat 2017;59:57–79. doi: 10.1111/anzs.12176.

31. Segata N, Izard J, Waldron L, et al. Metagenomic biomarker discovery and explanation. Genome Biol 2011;12:R60. doi: 10.1186/gb-2011-12-6-r60.

32. Breiman L; Random Forests. Machine Learning 2001;45(1):5–32. doi: 10.1023/A:1010933404324.

33. Liu C, Cui Y, Li X, et al. microeco: an R package for data mining in microbial community ecology. FEMS Microbiol Ecol 2020;97(2). doi: 10.1093/femsec/fiaa255.

34. Liaw A, Wiener M. Classification and regression by randomForest. R news 2002;2(3):18–22.

35. Holmes I, Harris K, Quince C. Dirichlet Multinomial Mixtures: generative models for microbial metagenomics. Plos ONE 2012;7:e30126. doi: 10.1371/journal.pone.0030126.

36. Moon KR, van Dijk D, Wang Z, et al. Visualizing structure and transitions in high-dimensional biological data. Nat Biotechnol 2019;37:1482–1492. doi: 10.1038/s41587-019-0336-3.

37. Zalar P, Gostinčar C, de Hoog GS, et al. Redefinition of Aureobasidium pullulans and its varieties. Stud Mycol 2008;61:21–38. doi: 10.3114/sim.2008.61.02.

38. Liu D, Chen Q, Zhang P, et al. The fungal microbiome is an important component of vineyard ecosystems and correlates with regional distinctiveness of wine. mSphere 2020;5. doi: doi:10.1128/msphere.00534-20.

39. Liu D, Howell K. Community succession of the grapevine fungal microbiome in the annual growth cycle. Environ Microbiol 2021;23:1842–1857. doi: 10.1111/1462-2920.15172.

40. Bauer F, Pretorius IS. Yeast stress response and fermentation efficiency: how to survive the making of wine. 2000.

41. Ciani M, Capece A, Comitini F, et al. Yeast interactions in inoculated wine fermentation. Front Microbiol 2016;7. doi: 10.3389/fmicb.2016.00555.

42. Costea PI, Hildebrand F, Arumugam M, et al. Enterotypes in the landscape of gut microbial community composition. Nat Microbiol 2018;3:8–16. doi: 10.1038/s41564-017-0072-8.

43. Hawkins DL, Ryder J, Lee SA, et al. Mixed yeast communities contribute to regionally distinct wine attributes. FEMS Yeast Res 2023;23. doi: 10.1093/femsyr/foad005.

44. Tempère S, Marchal A, Barbe J-C, et al. The complexity of wine: clarifying the role of microorganisms. Appl Microbiol Biotechnol 2018;102:3995–4007. doi: 10.1007/s00253-018-8914-8.

45. de Souza RSC, Okura VK, Armanhi JSL, et al. Unlocking the bacterial and fungal communities assemblages of sugarcane microbiome. Sci Rep 2016;6:28774. doi: 10.1038/srep28774.

46. Saunders AM, Albertsen M, Vollertsen J, et al. The activated sludge ecosystem contains a core community of abundant organisms. ISME J 2016;10:11–20. doi: 10.1038/ismej.2015.117.

47. Edwards J, Johnson C, Santos-Medellín C, et al. Structure, variation, and assembly of the root-associated microbiomes of rice. Proc Natl Acad Sci USA 2015;112:E911–E920. doi: doi:10.1073/pnas.1414592112.

48. Fitzpatrick CR, Copeland J, Wang PW, et al. Assembly and ecological function of the root microbiome across angiosperm plant species. Proc Natl Acad Sci USA 2018;115:E1157–E1165. doi: doi:10.1073/pnas.1717617115.

49. Kembel SW, O’Connor TK, Arnold HK, et al. Relationships between phyllosphere bacterial communities and plant functional traits in a neotropical forest. Proc Natl Acad Sci USA 2014;111:13715–13720. doi: doi:10.1073/pnas.1216057111.

50. Abdelfattah A, Freilich S, Bartuv R, et al.; Global analysis of the apple fruit microbiome: are all apples the same? Environmental Microbiology 2021;23(10):6038–6055. doi: 10.1111/1462-2920.15469.

51. Morrison-Whittle P, Lee SA, Goddard MR. Fungal communities are differentially affected by conventional and biodynamic agricultural management approaches in vineyard ecosystems. Agr Ecosyst Environ 2017;246:306–313. doi: 10.1016/j.agee.2017.05.022.

52. Onetto CA, Schmidt SA, Roach MJ, et al. Comparative genome analysis proposes three new Aureobasidium species isolated from grape juice. FEMS Yeast Res 2020;20. doi: 10.1093/femsyr/foaa052.

53. Rathnayake RMSP, Savocchia S, Schmidtke LM, et al. Characterisation of Aureobasidium pullulans isolates from Vitis vinifera and potential biocontrol activity for the management of bitter rot of grapes. Eur J Plant Pathol 2018;151:593–611. doi: 10.1007/s10658-017-1397-0.

54. Grube M, Schmid F, Berg G. Black fungi and associated bacterial communities in the phyllosphere of grapevine. Fung Biol 2011;115:978–986. doi: 10.1016/j.funbio.2011.04.004.

55. Prakitchaiwattana CJ, Fleet GH, Heard GM. Application and evaluation of denaturing gradient gel electrophoresis to analyse the yeast ecology of wine grapes. FEMS Yeast Res 2004;4:865–877. doi: 10.1016/j.femsyr.2004.05.004.

56. Castoria R, De Curtis F, Lima G, et al. Aureobasidium pullulans (LS-30) an antagonist of postharvest pathogens of fruits: study on its modes of action. Postharvest Biol Tech 2001;22:7–17. doi: 10.1016/S0925-5214(00)00186-1.

57. Sharma RR, Singh D, Singh R. Biological control of postharvest diseases of fruits and vegetables by microbial antagonists: A review. Biol Control 2009;50:205–221. doi: 10.1016/j.biocontrol.2009.05.001.

58. Briceño EX, Latorre BA. Characterization of Cladosporium Rot in grapevines, a problem of growing importance in Chile. Plant Dis 2008;92:1635–1642. doi: 10.1094/pdis-92-12-1635.

59. Nam MH, Park MS, Kim HS, et al. Cladosporium cladosporioides and C. tenuissimum cause blossom blight in strawberry in Korea. Mycobiology 2015;43:354–359. doi: 10.5941/MYCO.2015.43.3.354.

60. Lutz MC, Sosa MC, Colodner AD. Effect of pre and postharvest application of fungicides on postharvest decay of Bosc pear caused by Alternaria—Cladosporium complex in North Patagonia, Argentina. Sci Hortic 2017;225:810–817. doi: 10.1016/j.scienta.2017.05.007.

61. Albertin W, Setati ME, Miot-Sertier C, et al. Hanseniaspora uvarum from winemaking environments show spatial and temporal genetic clustering. Front Microbiol 2016;6. doi: 10.3389/fmicb.2015.01569.

62. Steenwyk JL, Opulente DA, Kominek J, et al. Extensive loss of cell-cycle and DNA repair genes in an ancient lineage of bipolar budding yeasts. PLOS Biol 2019;17:e3000255. doi: 10.1371/journal.pbio.3000255.

63. Haase MAB, Steenwyk JL, Boeke JD. Gene loss and cis-regulatory novelty shaped core histone gene evolution in the apiculate yeast Hanseniaspora uvarum. bioRxiv 2023:2023.08.28.551515. doi: 10.1101/2023.08.28.551515.

64. Torija MJ, Rozès N, Poblet M, et al. Yeast population dynamics in spontaneous fermentations: Comparison between two different wine-producing areas over a period of three years. A Van Leeuw J Microb 2001;79:345–352. doi: 10.1023/A:1012027718701.

65. Vigentini I, De Lorenzis G, Fabrizio V, et al. The vintage effect overcomes the terroir effect: a three year survey on the wine yeast biodiversity in Franciacorta and Oltrepò Pavese, two northern Italian vine-growing areas. Microbiol 2015;161:362–373. doi: 10.1099/mic.0.000004.

66. Evans KJ, Pirie AJG. Weather variables for within-vineyard awareness of Botrytis risk. Aust J Grape Wine R 2024;2024:6630039. doi: 10.1155/2024/6630039.

67. Sosnowski MR, Ayres MR, McCarthy MG, et al. Winegrape cultivars (Vitis vinifera) vary in susceptibility to the grapevine trunk pathogens Eutypa lata and Diplodia seriata. Aust J Grape Wine R 2021;28:166–174. doi: 10.1111/ajgw.12531.

68. Varela C. The impact of non-Saccharomyces yeasts in the production of alcoholic beverages. Appl Microbiol Biotechnol 2016;100:9861–9874. doi: 10.1007/s00253-016-7941-6.

