## Supplementary material for "Temporal and spatial dynamics within the fungal microbiome of grape fermentation": Fig. S1, Fig. S2, Fig. S3, Fig. S4

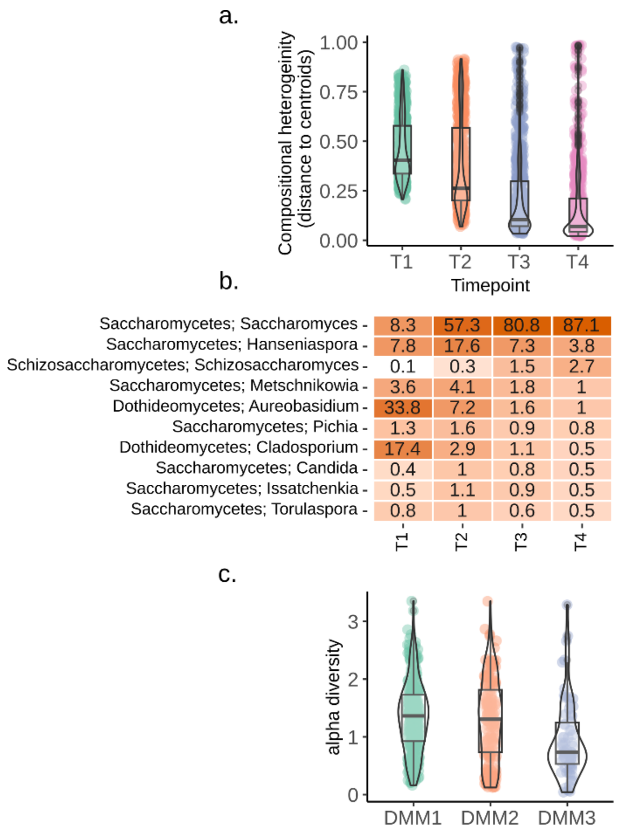


**Fig. S1. a** Compositional heterogeneity (beta-dispersion of Bray-Curtis distances) of samples separated by fermentation timepoint. **b** Relative abundance for the top 10 Genera separated by fermentation timepoint. **c** Alpha-diversity (Shannon index) of T2 samples belonging to each DMM partition.

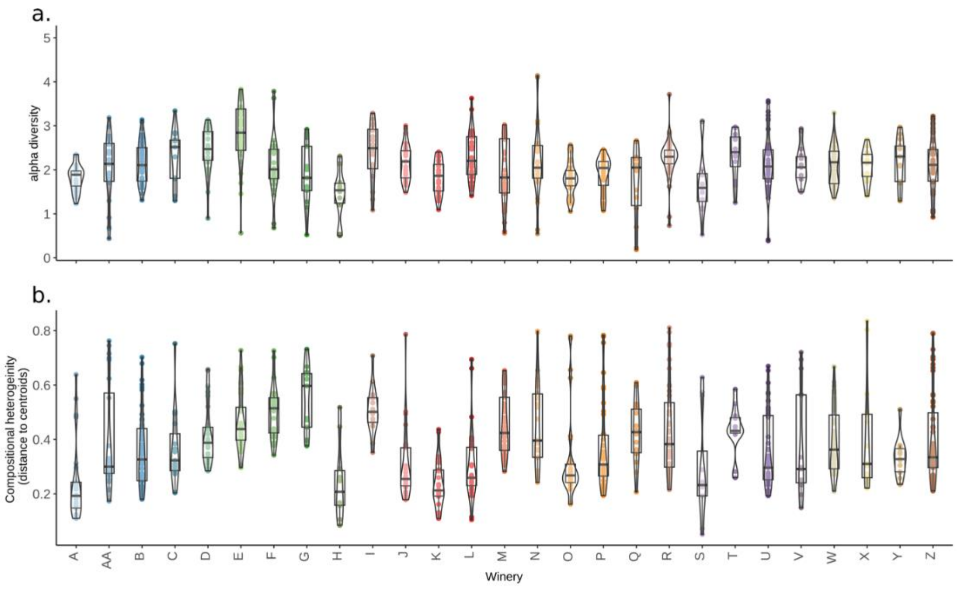


**Fig. S2**. **a** Violin-plot showing the alpha-diversity (Shannon index) and **b** compositional heterogeneity (beta-dispersion of Bray-Curtis distances) of T1 samples separated by winery. Statistical differences between regions are available in Supp Tables 2 and 4.

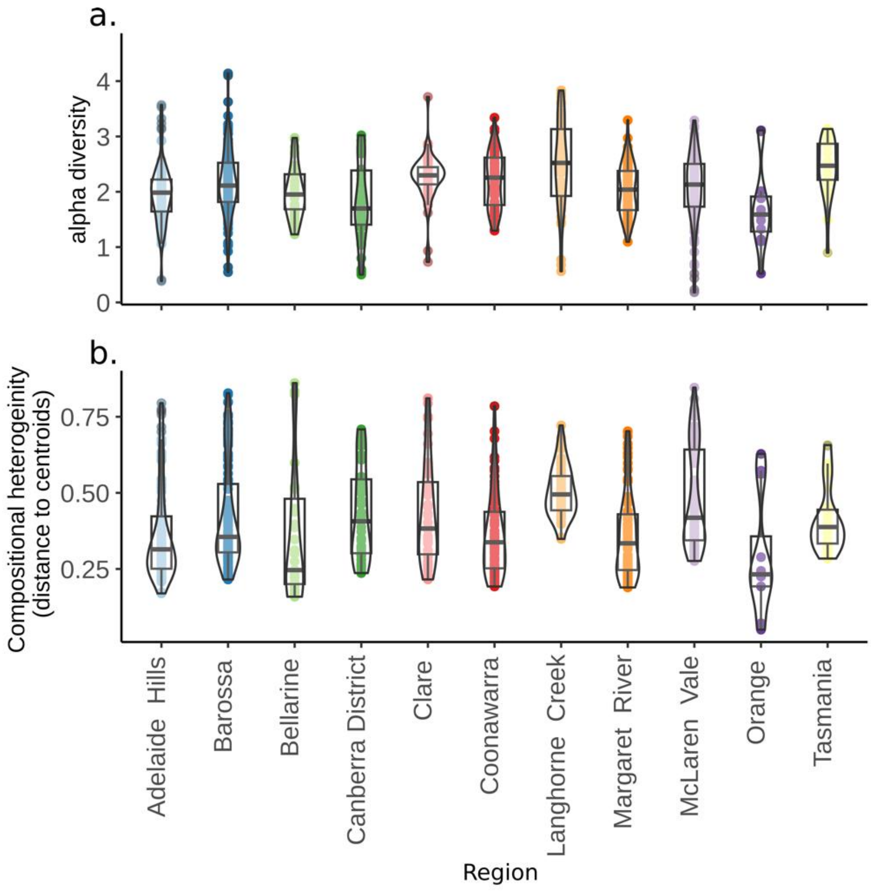


**Fig. S3**. **a** Violin-plot showing the alpha-diversity (Shannon index) and **b** compositional heterogeneity (beta-dispersion of Bray-Curtis distances) of T1 samples separated by wine-growing region. Statistical differences between regions are available in Supp Tables 3 and 5.

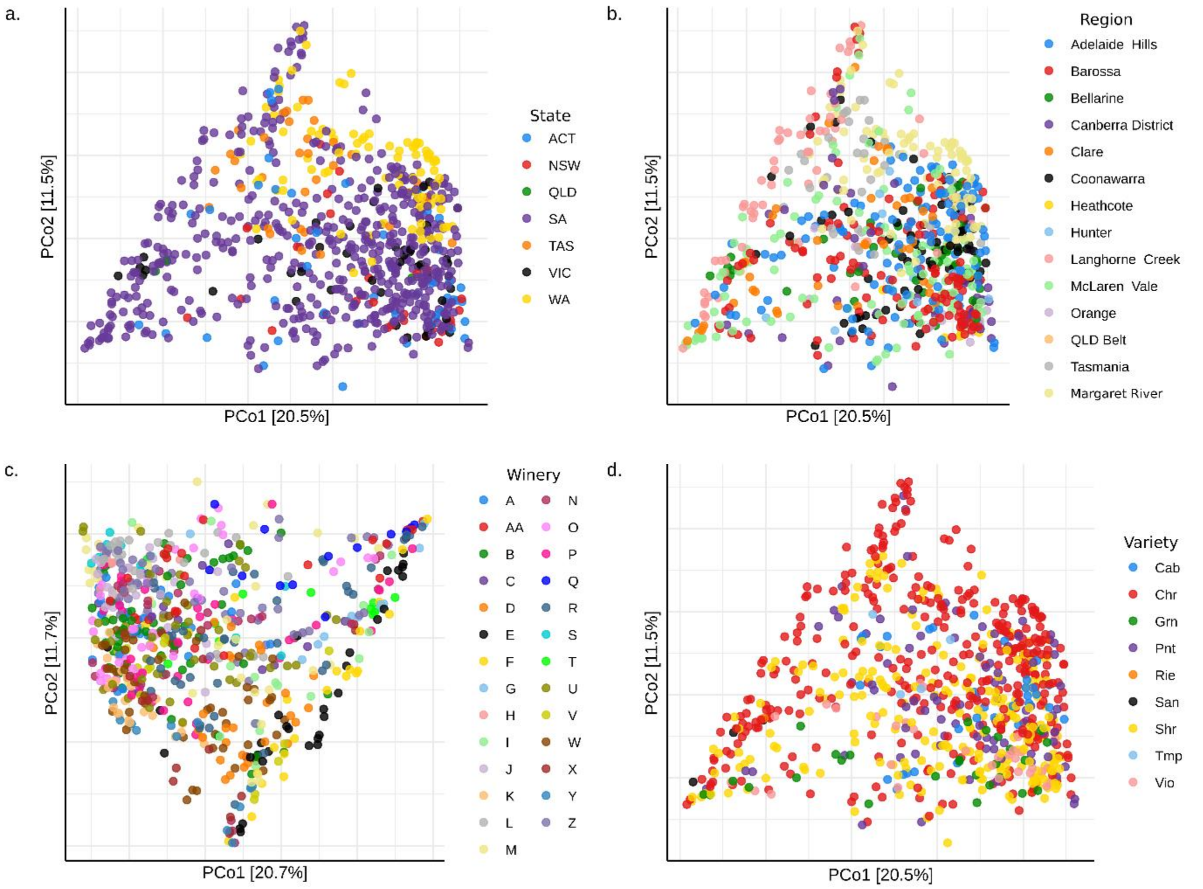


**Fig S4**. Principal coordinate analysis (PCoA) of T1 samples based on Bray-Curtis (BC) dissimilarities labelled by **a** state, **b** wine region, **c** winery and **d** grape variety. Only wineries with > 10 samples in total were included in the winery plot.
